# High Quality, Long-Read Genome Assemblies of Two Virulent Field Isolates of *Pyricularia oryzae* from Portugal

**DOI:** 10.64898/2026.06.12.731836

**Authors:** Pedro Rosa, João Bilro, Ricardo S. Ramiro, Cristina Azevedo

## Abstract

The fungal pathogen *Pyricularia oryzae* is notorious for causing blast disease in various important cereal crops, including wheat, rice, millet, and oat. Whole-genome-informed data on this pathogen are necessary to better understand the host adaptability of the fungus, including identifying key determinants of infection to enable more precise disease control. Here, we report highly contiguous genome sequences (using long-read PacBio technology) of two isolates from rice paddies in Portugal, M22.7 and T22.2, which exhibit distinctly aggressive symptoms in rice. Both mitochondrial and nuclear sequences were characterised in this study. The resulting nuclear genomes have assembly lengths of 46.4 Mb for M22.7 (198x coverage) and 46.3 Mb for T22.2 (163x coverage), with near-complete BUSCO completeness (98.8%) and a 0% contamination score (EukCC). Phenotypic analyses on infected rice leaves show that both isolates are infectious and virulent in two different rice cultivars. Most notably, M22.7 lacks the avirulence effector genes Avr-Pii, Avr-Pia and Avr-Pik, which are present in T22.2. This announcement represents the first genome resource for natural isolates of P. oryzae from Portugal in over 30 years, filling an important data gap from a major European rice-producing country that produces locally adapted rice varieties under specific agro-environmental conditions (near the Atlantic coast).

## Genome Announcement

*Pyricularia oryzae* (syn. *Magnaporthe oryzae*) is a hemibiotrophic plant pathogen, well studied for its infectivity in several cereal and grass species (Kato, 2001; Couch et al., 2005; Fernandez and Orth, 2018). It is primarily known for causing blast disease in rice and wheat, infecting all aboveground plant parts. In fields with low resistance pressure, *P. oryzae* can rapidly proliferate and cause severe and annual losses in rice production, in some cases leading to an almost complete loss of production regionally (Skamnioti and Gurr, 2009; Dean et al., 2012, Savary et al., 2019). As of July 2026, there were 582 genome assemblies of *P. oryzae* available on NCBI. The increasing availability of these genomic resources over the decades, along with the existence of reference rice genomes, has enabled a better understanding of the evolution of *P. oryzae* and of the selective pressures influencing the variability of both effector (*AVR*s) and plant resistance genes, respectively (Gallet et al., 2016; Yoshida et al., 2016; Wang et al., 2017). Phylogenetic inferences have also captured the evolutionary history of host-shift events and the subdivision of genetically distinct, host-defined lineages (Gladieux et al., 2018; Langner et al., 2018; Latorre et al., 2020). Portugal is the main rice consumer and one of the main producers in Europe (Fraga et al., 2019). Rice production is concentrated on locally adapted varieties (e.g. the Carolino variety (Japonica)) in Atlantic coastal estuaries under specific agro-environmental conditions, including multiple stresses such as drought, temperature extremes and soil salinity (Fraga et al., 2019). Rice blast causes severe and recurrent yield losses annually, and so, characterising *P. oryzae* genomes from Portugal is critical for both national rice production and effective global genomic surveillance. Several Portuguese *P. oryzae* field isolates were isolated in the early 1990s (Roumen et al., 1997), from which the only two publicly available genomic sequences of Portuguese isolates originated: isolates PR72, assembly acc. no. GCA_000734785.1; and PR0009, SRA acc. no. SRR6384771, assembly acc. no. DBPUGO000000000.1; however, no subsequent *P. oryzae* isolates from Portugal have been isolated and sequenced since then. In addition, as of July 2026, only 12 genomes isolated from rice in Europe had been deposited in NCBI, and all of them were isolated over 15 years ago. Consequently, we aimed to isolate and characterise novel Portuguese *P. oryzae* isolates through long-read sequencing, especially because no *P. oryzae* genomic resources from Portugal are available using these approaches. In addition to filling an important gap in the genomic characterisation of Portuguese *P. oryzae* populations, these resources will provide valuable information for the national rice breeding programme, in which resistance to *P. oryzae* is an important breeding target. These novel genomic resources provide a genetic framework to better understand the diversity and virulence of the pathogen population against which new rice varieties will be deployed. By linking pathogen genotypes with observed infection phenotypes, the programme can select representative and relevant isolates for resistance screening, identify potential vulnerabilities of resistance genes, and also support the identification and combination of resistance sources with wider effect. This supports the predictive value of breeding trials and reduces the risk of selecting varieties whose resistance is effective only against a limited number of the local pathogen population. As such, between 2022 and 2024, *P. oryzae* isolates were collected from the three main rice-growing regions of Portugal: Mondego, Tejo, and Sado. Two isolates, termed M22.7 and T22.2, were isolated during early to mid-August 2022 from blast-infected samples collected in the Mondego and Tejo River basins, respectively, and have been characterised.

Isolate T22.2 was obtained from infected parts of a cultivated rice variety (var. Lusitano), and M22.7 from an infected wild rice sample (*Oryza* sp., Fig. 1a). Information regarding the collection and isolation source of these isolates is shown in Table 1. Small cuts of unsterilised pieces of leaves (1-2 cm^2^), panicle necks or whole single grains showing blast lesions were placed on 2% agar (w/v) and incubated at 24-25 °C under constant light for at least 48 hours. Single spores or *P. oryzae-*like hyphal growth were isolated using a sterile syringe or scalpel, respectively, transferred to complete medium (Talbot et al., 1993) and incubated at 24-25 °C in a 12h/12h photoperiod for at least 48 hours, monitoring growth and possible contaminants. Pure isolate cultures were confirmed as *P. oryzae* by running a PCR against the *Pyricularia*-specific *Pot2* fragment (Harmon et al., 2003), sequencing the internal transcribed spacer ITS1-ITS4 marker region (Fig. 1b) and morphological comparisons with reference lab strains (Fig. 1c).

**Figure 1.**
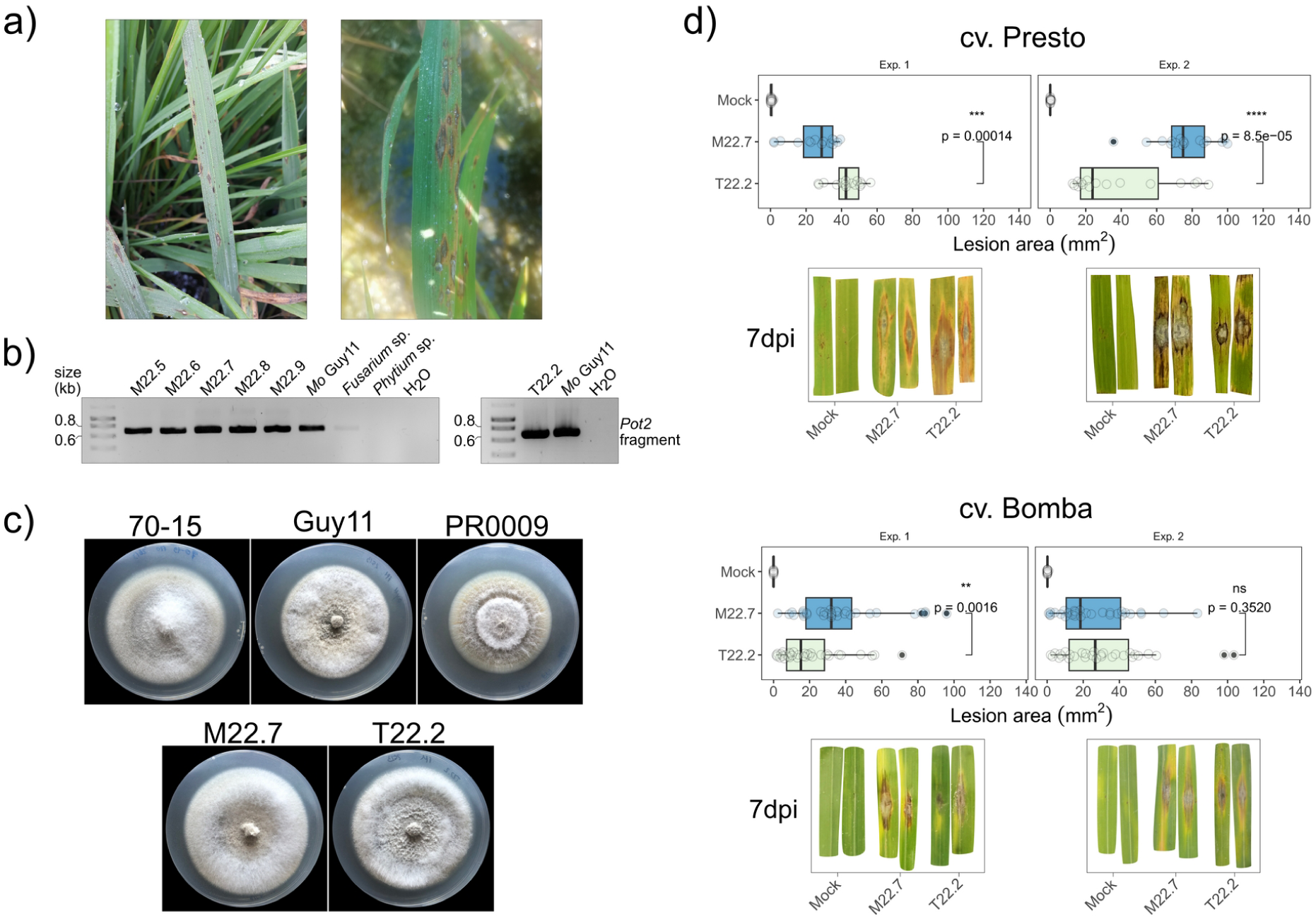
– Differences between *P. oryzae* isolates M22.7 and T22.2. a) Cultivated rice plants at early August 2022 showing blast symptoms in two rice paddies from Mondego (left) and Tejo (right). b) Diagnostic PCR detecting both M22.7 and T22.2 as *Pot2*-positive; *Mo* Guy11 was used as a positive control, and *Fusarium* sp. and *Phytium* sp. as negative controls. c) Ten-day-old culture plates of *P. oryzae* lab strains 70-15 and Guy11 and of Portuguese field isolates in complete medium (note the existence of denser aerial hyphae in M22.7 compared with T22.2) in Complete Media. d) Detached rice leaves of cv. Presto (left) and cv. Bomba (right) prick-inoculated with M22.7 and T22.2 conidia (at 10^5^ conidia/mL in 0.02% Tween-20 + 0.25% gelatin from porcine skin). Symptoms were quantified as lesion area (mm^2^) at 7 days post-infection (dpi). Statistical differences between isolate-specific infection and respective p-values shown in each plot (One-way ANOVA and post-hoc Tukey test, cv. Presto infection: n = 15 and n = 16 and respectively in experiments 1 and 2; cv. Bomba infection: n = 32 and n = 30 respectively in experiments 1 and 2).

**Table 1.** – Summary of the collection/isolation information of *P. oryzae* ‘M22.7’ and ‘T22.2’ isolates.

| Parameter | Value | *P. oryzae* 'M22.7' | *P. oryzae* 'T22.2' |
| --- | --- | --- | --- |
| Collection | Date | 2<sup>nd</sup> August 2022 | 18<sup>th</sup> August 2022 |
| Geolocation | Country | Portugal | Portugal |
|  | Region | Montemor-o-Velho | Coruche |
|  | Latitude | 40.1355 N | 38.9492 N |
|  | Longitude | 8.6687 W | 8.5123 W |
| Isolation | Source | rice-producing basin | rice-producing basin |
|  | Hydrographic basin | Mondego river | Tejo river |
|  | Material | leaves, panicles and grains | leaves, panicles and grains |
|  | Host | *Oryza sativa* | *Oryza sativa* |
|  | Host variety | wild rice | Lusitano |
| Other | Strain | wild-type | wild-type |
|  | Mating type | Mat1-1 | Mat1-1 |

Both isolates were determined to be virulent in the susceptible rice cultivars Presto and Bomba *ex vivo* (Fig. 1d). Both isolates were grown in a rice flour agar medium (Le Naour-Vernet et al., 2023) under constant light for at least 7 days to induce conidiation. Then, well-established leaves were cut from the plants at the 2-4 leaf stage into roughly 4 cm segments, placed adaxial side-up in 0.6% (w/v) plant agar plates supplemented with 3 µg/mL 6-benzylaminopurine, and 4 µL of a conidial suspension at 1×10^5^ conidia/mL (in 0.02% Tween-20 + 0.25% gelatine from porcine skin) was spotted onto each leaf segment, followed by wounding it at the inoculation point using a sterile 21G hypodermic needle. Symptoms were assessed 7 days post-infection by measuring the area of necrotic and chlorotic tissue surrounding the pricked inoculation point using ImageJ v1.54g20 (Fiji).

Finally, genomic DNA was extracted and purified from these two isolates using a hexadecyltrimethylammonium bromide extraction procedure. Purity of DNA samples was measured using a μDrop plate (Thermo Fisher, USA), and quantification was performed using a Quantus Fluorometer (Promega, USA). Library preparation and HiFi sequencing were outsourced to Edinburgh Genomics (The University of Edinburgh, Scotland). Briefly, PacBio SMRTbell libraries were constructed with a minimum of 5 µg of DNA using the PacBio SMRTBell prep kit 3.0 (PacBio, USA) and loaded onto the PacBio Sequel IIe sequencer (HiFi mode) using the PacBio Sequel II Binding kit 3.2. Sequencing generated 585,514 reads with an average size of 16,724 bp for isolate M22.7 and 423,346 reads with an average size of 18,356 bp for T22.2 (Table 2).

**Table 2.** – Summary of the statistics for the final assembly and annotation of *P. oryzae* ‘M22.7’ and ‘T22.2’ isolates.

| Category | Metric | *P. oryzae* 'M22.7' | *P. oryzae* 'T22.2' |
| --- | --- | --- | --- |
| Sequencing statistics | Total read count | 585514 | 423346 |
|  | Total read length (bp) | 9792204854 | 7770920304 |
|  | Median read length (bp) | 16229 | 17693 |
|  | Longest read length (bp) | 44156 | 50727 |
| Assembly statistics | Preferred assembler | NextDenovo | NextDenovo |
|  | Total assembly length (Mb) | 46.4 | 46.3 |
|  | Number of contigs | 26 | 15 |
|  | Largest contig length (bp) | 8238573 | 9080260 |
|  | GC content | 50.87% | 51.02% |
|  | N50 | 4729120 | 6522540 |
|  | N90 | 1203062 | 3967384 |
|  | L50 | 4 | 4 |
|  | L90 | 10 | 7 |
|  | Coverage depth | 198x | 163x |
| Completeness (BUSCO) | BUSCO completeness | 98.8% | 98.8% |
|  | Complete BUSCOs | 4439 | 4440 |
|  | Single-copy BUSCOs | 4432 | 4433 |
|  | Duplicated BUSCOs | 7 | 7 |
|  | Fragmented BUSCOs | 19 | 16 |
|  | Missing BUSCOs | 34 | 36 |
| Contamination (EukCC) | Completeness score | 100.0% | 100.0% |
|  | Contamination score | 0.0% | 0.0% |
|  | Maximum silent contamination | 11.79% | 1.37% |
| Annotation statistics | Total predicted gene models | 12731 | 13065 |
|  | mRNAs | 12402 | 12733 |
|  | tRNAs | 329 | 332 |
|  | GO terms | 7042 | 7140 |
|  | InterPro terms (InterProScan) | 8800 | 8943 |
|  | EggNOG terms (EggNOG-mapper) | 10516 | 10720 |
|  | Pfam domains | 7647 | 7781 |
|  | CAZymes (dbCAN3) | 537 | 553 |
|  | Proteases (MEROPS) | 378 | 385 |
|  | KEGG orthology (kofamscan) | 4265 | 4273 |
|  | Biosynthetic gene clusters (antiSMASH) | 63 | 67 |
|  | Predicted secreted genes | 1688 | 1765 |
|  | Predicted effectors (EffectorP) | 631 | 664 |
| Mitochondrial statistics | Total assembly length (bp) | 34866 | 34866 |
|  | Number of contigs | 1 | 1 |
|  | Coverage depth | 6520x | 5605x |
|  | Predicted mitochondrial genes | 44 | 44 |

The sequenced reads from both isolates were assembled *de novo* using hifiasm v0.25.0-r726 (Cheng et al., 2021) and NextDenovo v2.5.2 (Hu et al., 2024). The former was executed using default parameters, and the latter was executed with slight modifications to accommodate the correct assembly of the raw PacBio HiFi reads (read_type = hifi, genome_size = 41m, pa_correction = 2, sort_options = -m 450g -t 64). Quality control metrics, such as relative GC content, N50, N90, and others, were calculated for both resulting assemblies using Blobtoolkit v4.4.5 (Challis et al., 2020). Completeness was assessed with BUSCO v5.8.3 (Manni et al., 2021), based on evidence from the lineage-specific dataset ‘sordariomycetes_odb12’ (Tegenfeldt et al., 2025). Relative contamination of the assemblies was determined using EukCC v2.1.2 (Saary et al., 2020). Both NextDenovo assemblies were of high quality, showing the same completeness/contamination scores: BUSCO completeness of 98.8%, EukCC completeness of 100%, and a contamination score of 0% (Fig. 2A and Table 2). Based on these metrics, the NextDenovo assemblies for both isolates were identified as the most contiguous and complete and were kept as the final assemblies. The raw reads were aligned to the assembled genomes to calculate genome coverage with minimap2 v2.27-r1193. Then, the assemblies were filtered based on taxonomic classification obtained by mapping all contigs against NCBI’s BLAST v5 ‘nt/nr’ database (release 2025_08) and UniProt’s reference genomes (release 2025_03) using BLAST v2.12.0 ‘blastn’ (Camacho et al., 2009) and DIAMOND v2.1.13 ‘blastx’ searches (Buchfink et al., 2021), respectively. The results were aggregated and consolidated following a majority rule to select the best hits for each contig. This removed one mitochondrial contig assigned to a *Penicillium* spp. in the M22.7 assembly, while the T22.2 assembly remained unaltered. Because *Penicillium* is a common contaminant in *P. oryzae* cultures, we do not exclude the possibility that the presence of this contig could be due to an overlooked, isolated biological contamination in M22.7 cultures (Molinari and Talbot, 2022). Notably, a single mitochondrial contig was identified for each assembly. The mitochondrial genome of each isolate was assembled and annotated with MitoHiFi v3.2 (Laslett and Canbäck, 2008; Allio et al., 2020; Uliano-Silva et al., 2023), using the mitochondrial genome of strain Guy11 as reference (GenBank OP095391.1, (Xu et al., 2023)). This resulted in the assembly of the complete mitochondrial genome for each isolate, spanning 34866 bp, and containing 44 annotated genes, corresponding to a single mitochondrial contig from each assembly. After removing these contigs, the total length of the final genome assemblies was 46.4 Mb (26 contigs, N50 of 4.73 Mb) for M22.7 and 46.3 Mb (15 contigs, N50 of 6.52 Mb) for T22.2. Lastly, the quality of the final genome assemblies was evaluated with Inspector v1.3.1 (Guo et al., 2025) based on its computed quality values (QV) and the identification of structural errors. For this purpose, minimap2 v2.31-r1302 was used to align the raw reads back to the assemblies and identify misassemblies and structural errors. M22.7 had a QV of 54.6 and an 85 bp collapse in contig ctg000080 and T22.2 had a QV of 61.9 with no structural errors. QUAST v5.3.0 (Mikheenko et al., 2023) was also used to assess genome assembly contiguity and quality based on comparisons against the reference genome assembly of the rice-infecting isolate 70-15 (assembly acc. no. GCA_000002495.2). QUAST results showed high contiguity and a low rate of structural discrepancies between both NextDenovo assemblies and the reference genome. In fact, for both comparisons against 70-15, only ∼0.12% of the assembled bases were estimated to be misassembled. Along with previously computed BUSCO completeness values and EukCC contamination scores, these results further demonstrate the high quality of both final genome assemblies. Table 2 shows assembly statistics.

**Figure 2.**
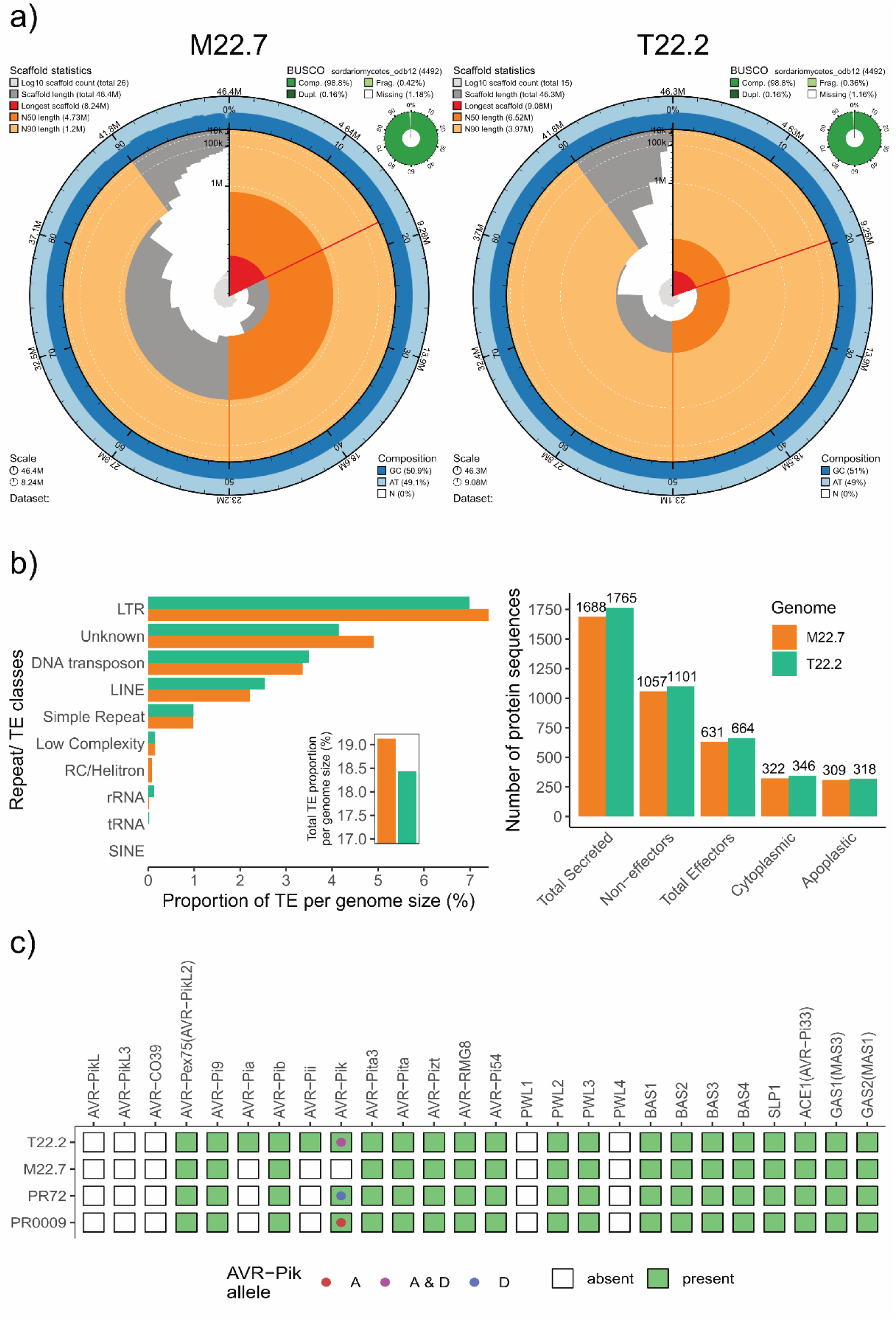
– Genomic differences between assembled genomes of M22.7 and T22.2. a) Snail plots from final assemblies (NextDenovo) of M22.7 and T22.2, computed with Blobtoolkit v4.4.5. Several scaffold statistics and metrics, BUSCO completeness values, and GC composition percentages are shown for each sequenced isolate. b) Left, proportion of different transposable element (TE) classes identified by RepeatMasker by assembly length (inset: total proportion of TEs for each assembly); right, number of putative secreted and cytoplasmic or apoplastic effector genes identified by EffectorP from sequenced isolates. c) Effector presence/absence and allelic variation of *AVR-Pik* between sequenced Portuguese *P. oryzae* isolates.

To place the genome assemblies in a phylogenetic context, a phylogeny was inferred for a subset of representative *P. oryzae* genomes (see Supplementary Figure 1 and its legend). The entire topology was highly supported (UFBoot > 94%), and isolates M22.7 and T22.2 formed a fully supported separate branch stemming from a shared ancestor with PR0009 (Supplementary Figure 1). Additionally, PR72 and FR13 formed a separate sister branch, with both branches belonging to a clade solely comprised of rice-infecting isolates, as expected.

Pairwise structural genome comparisons were computed to assess assembly structure and contiguity, using minimap2 v2.24 through D-GENIES v1.5.0 (Cabanettes and Klopp, 2018), with the ‘Many repeats’ option. We compared both isolates against each other, and each isolate against three reference rice-infecting genomes (70-15, FR13 and Guy11). The dotplots show high contiguity between our isolates and in comparison to the reference genomes (Supplementary Figures 2 and 3); but M22.7, in general, showed more structural differences (e.g. inversions, transpositions, etc.) than T22.2. The presence of mini-chromosomes, which contribute to the natural variation in *P. oryzae* genomes (Peng et al., 2019; Langner et al., 2021), was not assessed in this study and would require karyotype analysis by CHEF electrophoresis.

For genome annotation, gene models were predicted using the Funannotate v1.8.17 annotation pipeline (Jonathan M. Palmer and Jason Stajich, 2020). Protein and transcript evidence data were collected from the Mycocosm public database (https://mycocosm.jgi.doe.gov/mycocosm/home) (Grigoriev et al., 2014), namely from the comparative multigene cluster ‘MagorFR13_1 comparative clustering 4830’. The annotation process also included a search for secondary metabolite biosynthetic gene clusters (BGCs) using the fungal version of antiSMASH v7.1.0 (Blin et al., 2023), CAZymes from dbCAN3 v12.0 (Zheng et al., 2023), proteases from MEROPS v12.0 (Rawlings et al., 2018), transfer RNAs with tRNAscan-SE v2.0.12 (Chan et al., 2021) and COGs and GO terms with eggNOG-mapper v2.1.12 (Cantalapiedra et al., 2021). Pfam functional data were also integrated into the InterPro annotation (release 2025_04) using InterProScan v5.73-104.0 (Jones et al., 2014). Also, Kofamscan v1.3.0 (Aramaki et al., 2020) was used with mostly default parameters (--profile=eukaryote.hal --format=detail-tsv --e-value=0.01) to annotate the predicted gene models with the KEGG orthology (KO) groups based on the profile Hidden Markov Models (HMMs) list from the KOfam database (release 2025_05). This gene prediction approach revealed an average of 3.16 genes for M22.7 and 3.26 genes per 10kb window for T22.2. The isolate T22.2 was predicted to have 334 more genes than M22.7, with 13,065 and 12,731 predicted gene models, respectively, similar to the reference sequence of strain 70-15 (13,184 gene models). On average, over 80% of all predicted genes had at least one predicted function or domain based on the EggNOG, Pfam, and InterPro databases, and an average of 2.7% were identified as tRNAs. Repetitive sequences were predicted and identified with RepeatModeler v2.0.6 (Flynn et al., 2020) for both final genome assemblies. Then, RepeatMasker v4.1.8 (Smit, AFA, Hubley, R & Green, 2014) was used to mask the assemblies, extract the predicted repetitive elements and classify them into different families. Both assemblies showed a similar repeat content as identified by RepeatMasker (Fig. 2b, left), with approximately 19.1% in M22.7 assembly and 18.4% in T22.2, mainly simple repeats but also *Gypsy* LTR and long interspersed nuclear retrotransposon elements (LINEs) (Fig. 6a). Interestingly, M22.7 contains 24 insertions of *Helitron* transposable elements (TEs) whilst none were identified in T22.2. Finally, we sought to determine differences between the two isolates in their effector gene profiles. To predict the secretome, Phobius v1.01 (Käll et al., 2004) and SignalP v6.0 (Teufel et al., 2022) were used to identify predicted secreted proteins. Then, each isolate’s secretome was passed to EffectorP v3.0 (Sperschneider and Dodds, 2022) to predict fungal effectors, with default parameters. The effector gene sequence dataset from (Petit-Houdenot et al., 2020) was also used to search each assembly using nucleotide local alignments computed with blastn v2.16.0 (Altschul et al., 1990), with a sequence identity and query coverage percentage thresholds of 80%, and an e-value cutoff of 1e-20. Overall, 1765 secreted protein genes were predicted in T22.2 and 1688 in M22.7. Relative to predicted effector-coding genes, T22.2 is predicted to encode 318 and 346 putative apoplastic and cytoplasmic effectors, respectively, while M22.7 is predicted to encode 309 apoplastic and 322 cytoplasmic putative effector genes (Fig. 2b, right and Table 2). This difference in effector-coding genes could simply reflect the overall difference in gene number between the two sequenced isolates. A subsequent pangenomic analysis including these and other *P. oryzae* isolates, coupled with infection data, could help determine whether these differences in effector-coding gene profiles have functional implications for strain pathogenicity. Unexpectedly, M22.7 does not encode the avirulence genes *Avr-Pii*, *Avr-Pia* or *Avr-Pik*, whereas T22.2 does (Fig. 2c). We also confirmed the absence of Avr-Pik in M22.7 by PCR (Supplementary Figure 4) using conserved primers (Supplementary Table 1). In fact, both *Avr-PikA* and *Avr-PikD* variants are present in T22.2 and have also recently been discovered in mini-chromosomes from two different strains containing both alleles (Kusaba et al., 2014; Langner et al., 2021). Nonetheless, both M22.7 and T22.2 are similarly infectious in detached leaves of the Bomba and Presto cultivars (Fig. 1d). This suggests that loss of Avr-Pik is not associated with reduced virulence under the tested conditions. The absence of Avr-Pik is still noteworthy, as this gene typically does not exhibit presence/absence polymorphisms (Kanzaki et al., 2012, Oikawa et al., 2024). Whether the absence of Avr-Pik incurs in fitness costs in other genetic backgrounds, host cultivars, or ecological contexts remains to be determined. However, M22.7 shows a consistent, highly reduced conidiation phenotype *in vitro* relative to T22.2 when grown in rice flour agar or in 2% water agar under constant light (Supplementary Figure 5). Often, aerial hyphae of M22.7 grown on solid medium would have to be removed and cultures incubated at least one more day under constant light to obtain sufficient conidia for *ex vivo* infections. This could suggest a fitness disadvantage, although this phenotype would have to be determined as well *in planta* to confirm this difference.

The genomes and functional annotations provided here offer modern genetic resources for *P. oryzae* from Southern Europe, making them extremely valuable for rice blast monitoring and detection. The completeness and assembly contiguity achieved by HiFi sequencing technologies can enable high-confidence structural variation and comparative studies that improve understanding of the genetic and pathogenic variability of rice blast. These two isolates are also available in the certified MUM fungal collection (University of Minho).

## Acknowledgments

The authors acknowledge Nick Talbot (The Sainsbury Lab, Norwich, UK), Stefan Jacob (IBWF, Mainz, Germany) and Pierre Gladieux and Didier Tharreau (Plant Health Institute, Montpellier, France) for providing *P. oryzae* strains Guy11, 70-15 and isolate PR0009, respectively, and those who facilitated our access to the rice fields: António Jordão (CCDRC), Daniel Carvalheiro, Lourenço Palha (COTArroz), and Rodrigo Capela (Aparroz). The authors also acknowledge Célia Soares (CEB-Biological Engineering Centre, University of Minho, Portugal) for receiving and storing both isolates in the MUM fungal collection. The authors also acknowledge Edinburgh Genomics for DNA sequencing and technical support.

## Data availability

The genomes of the Pyricularia oryzae M22.7 and T22.2 isolates are available on the National Centre for Biotechnology Information (NCBI) under the BioProject accession PRJNA1451107. The corresponding BioSample accessions are SAMN57182087 (M22.7) and SAMN57183957 (T22.2). Genome assemblies have been deposited under GenBank accession numbers JBXBCC000000000 (M22.7) and JBXBCB000000000 (T22.2). Genome annotations were published for each respective isolate on the following Zenodo repository: https://zenodo.org/records/19556489. The annotated mitochondrial genomes of both isolates have also been deposited on NCBI under GenBank accession numbers PZ276150 and PZ276151, respectively.

