## Supplementary Materials for "High Quality, Long-Read Genome Assemblies of Two Virulent Field Isolates of *Pyricularia oryzae* from Portugal"

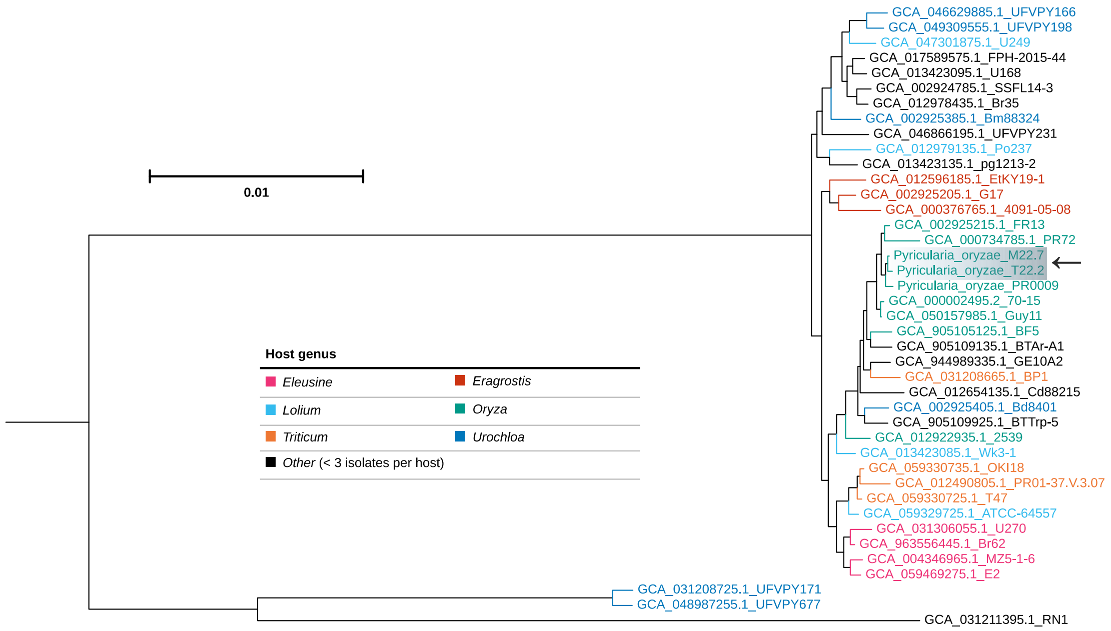


**Supplementary Figure 1** – Maximum-likelihood phylogeny of the representative genome assemblies of *Pyricularia oryzae*, including our sequenced isolates M22.7 and T22.2 (highlighted). Colours were attributed to each respective host plant genus that had at least 3 different isolates present in the phylogeny. In order to infer this phylogeny, which included our sequenced isolates and all of the genetic diversity of the *P. oryzae* genome assemblies present in NCBI as of July 2026, the required genomic data was gathered using NCBI Datasets v16.27.2; atypical or multi-isolate assemblies were excluded from this. We recovered 582 genome assemblies and computed pairwise average nucleotide identity (ANI) values using drep v3.6.2 with the fastANI algorithm at a 99.5% identity threshold. drep classified all assemblies into genetic clusters, and then chose a representative assembly per cluster (drep dereplicate). To this subset, we added three reference genome assemblies (FR13, Guy11 and 70-15, which were chosen to replace the representative assembly of their cluster due to their comparative relevance), two Portuguese isolate assemblies (PR72 and PR0009) and our isolates' final assemblies. We then assessed genome completeness for each assembly using BUSCO v5.8.3, with the 'sordariomycetes_odb12' reference dataset. We discarded assemblies with a BUSCO completeness value below 80%. This totaled to 42 genome assemblies and their BUSCO results. We then run BUSCO_Phylogenomics v20260630 (https://github.com/jamiemcg/BUSCO_phylogenomics), which generated an amino-acid supermatrix containing all complete BUSCOs present in at east 50% of the assemblies. The phylogeny was inferred from the supermatrix using IQTREE v3.1.3 with ModelFinder, 1000 ultrafast bootstrap (UFBoot) support replicates and 1000 replicates of the SH-like aLRT test (iqtree -s SUPERMATRIX.fasta --seed 42 --bnni -B 1000 --alrt 1000 -m TEST -nt 64). Tree annotations were carried out using TreeViewer v2.2.0.

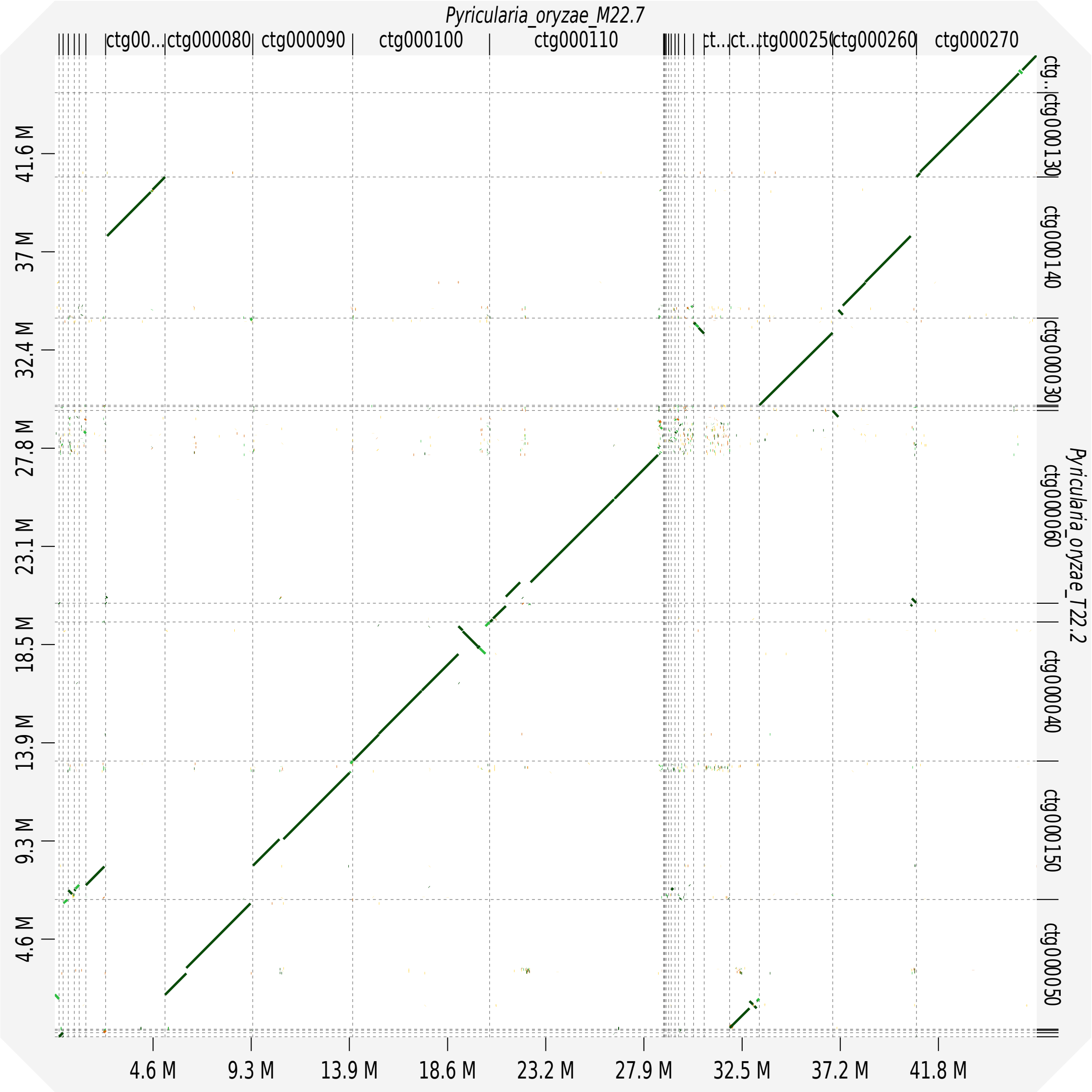


**Supplementary Figure 2** – Dotplot of the pairwise comparison of *Pyricularia oryzae* M22.7 and T22.2.

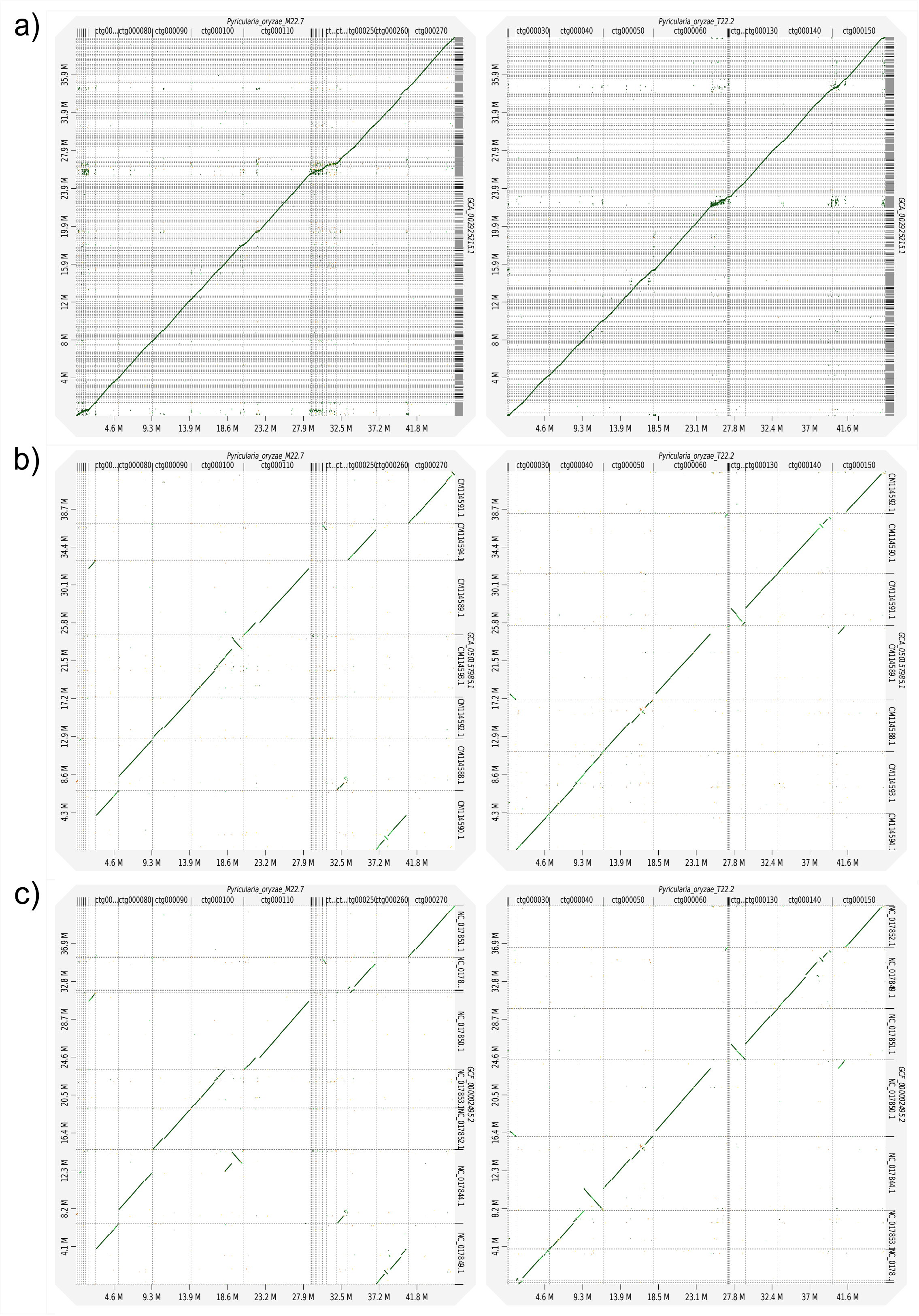


**Supplementary Figure 3** – Dotplots of pairwise comparisons between *Pyricularia oryzae* M22.7 (left) and T22.2 (right) with reference genome assemblies. a) Comparison with FR13; b) Comparison with Guy11; c) Comparison with 70-15.

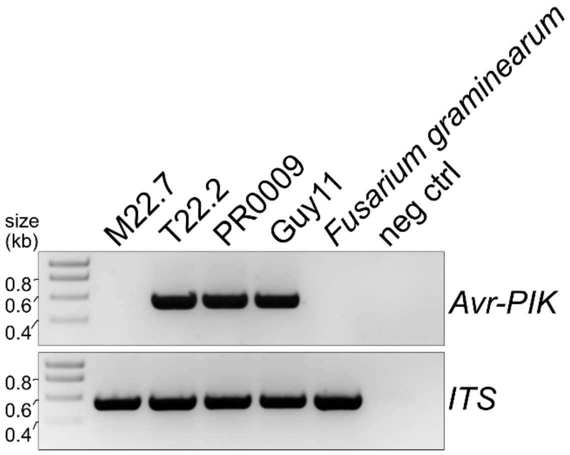


**Supplementary Figure 4** – Gel electrophoresis (1.2% agarose gel) detecting the presence/absence of *Avr-Pik* from *Pyricularia oryzae* isolates M22.7 and T22.2. Strain Guy11 was used as a positive control, and *Fusarium graminearum* as a negative control. Amplification of the ITS1-ITS4 marker region was run as an internal control.

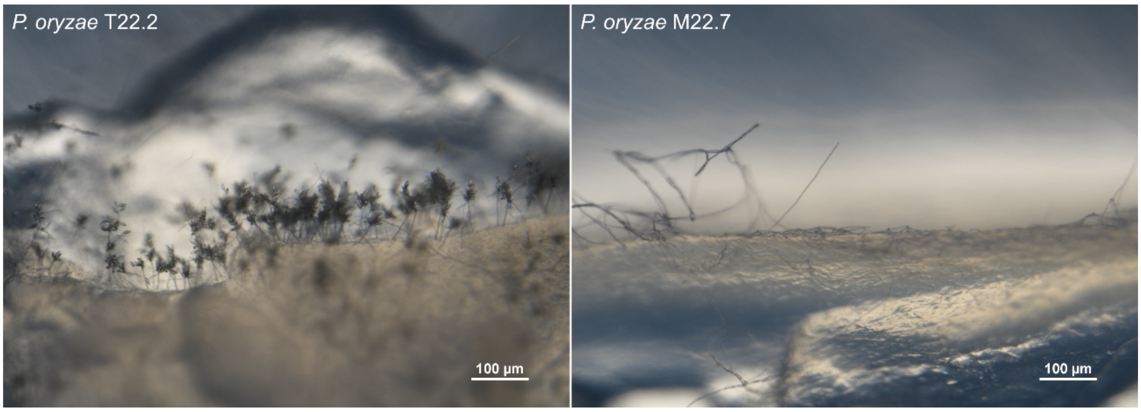


**Supplementary Figure 5** – *Pyricularia oryzae* conidiophore development on a 2% water agar (w/v) from isolates T22.2 (left) and M22.7 (right) at two days post-inoculation.

**Supplementary Table 1 –** Primers used in this study to detect the *Avr-Pik* locus

| **Primer name** | **Sequence (5'-3')** | **Amplicon size (bp)** | **Melting temperature (ºC)** |
|---|---|---|---|
| Forward primer* | cactctgcacacatcaacagtc | 535 bp | 58ºC |
| Reverse primer* | cttatgagccgtcaaccaagc | | 59ºC |

* Conserved flanks close to the *AVR-Pik* locus were searched among *Pyricularia oryzae* genomes through blastn using the wgs database from NCBI as well as in the T22.2 and M22.7 assemblies. Primers were designed to amplify the full-length *AVR-Pik* including 40-70bp flanks.
